# Erythrocyte-targeted immunomodulatory antigens enabled by in vivo selection of D-peptides

**DOI:** 10.1101/2020.07.15.203331

**Authors:** Alexander R. Loftis, Genwei Zhang, Coralie Backlund, Anthony J. Quartararo, Novalia Pishesha, Carly K. Schissel, Daniel Garafola, Andrei Loas, R. John Collier, Hidde Ploegh, Darrell J. Irvine, Bradley L. Pentelute

**Affiliations:** Department of Chemistry, Massachusetts Institute of Technology, Cambridge, MA 02139, USA; The Koch Institute for Integrative Cancer Research, Massachusetts Institute of Technology, Cambridge, MA 02142, USA; Program in Cellular and Molecular Medicine, Boston Children’s Hospital, Boston, MA 02115, USA; Broad Institute of MIT and Harvard, Cambridge, MA 02142, USA; Department of Microbiology, Harvard Medical School, Boston, MA 02115, USA; Department of Biological Engineering, Massachusetts Institute of Technology, Cambridge, MA 02139, USA; Ragon Institute of Massachusetts General Hospital, Massachusetts Institute of Technology and Harvard University, Cambridge, MA 02139, USA; Department of Materials Science and Engineering, Massachusetts Institute of Technology, Cambridge, MA 02139, USA; Howard Hughes Medical Institute, Chevy Chase, MD 20815, USA; Center for Environmental Health Sciences, Massachusetts Institute of Technology, Cambridge, MA 02139, USA

## Abstract

Targeting of antigens to erythrocytes can be used to selectively mitigate their immunogenicity, but the methods to equip a variety of cargoes with erythrocyte-targeting properties are limited. Here we identified a D-peptide that targets murine erythrocytes and decreases anti-drug antibody responses when conjugated to the protective antigen from *Bacillus anthracis*, a protein of therapeutic interest. The D-peptide likewise decreases inflammatory anti-ovalbumin (OVA) CD8^+^ T cell responses when attached to a peptide antigen derived from OVA. To discover this targeting ligand, we leveraged mass spectrometry to decode a randomized D-peptide library selected in mice, extending the application of synthetic libraries to in vivo affinity selections.

## Main

As the principal cellular component of blood, erythrocytes not only play a critical role in the transport of oxygen but are also implicated in the induction of antigen tolerance^1–4^. Covalent or non-covalent loading of antigens onto erythrocytes can drive antigen-specific tolerance, which is of great interest in mitigating formation of anti-drug antibodies and other adverse inflammatory and autoimmune reactions. While promising, erythrocyte antigen loading strategies are often complex, requiring ex vivo manipulation or bioconjugation to a large antibody.

Cell-targeting peptides (CTPs) provide an attractive alternative targeting strategy, as protein and peptide antigens can be easily attached to target-specific CTPs^5,6^. However, CTPs are typically composed of L-amino acids (L-peptides or L-CTPs), and therefore have had limited clinical impact due to their susceptibility to proteolysis.

One method to reduce susceptibility to proteolysis of L-CTPs is to use D-amino acids instead. For example, L-CTPs found in nature or from biological libraries (e.g., phage display) can be synthesized as a “retro-inverso” form to yield the D-enantiomer with inverted amino acid sequence^7–11^. In these cases, retention of binding affinity is difficult to predict. Alternatively, a one-bead-one-compound (OBOC) screening approach can be used to discover novel D-enantiomer CTPs, but this requires covalent attachment of each library member to a solid support^12^. Because of this limitation, OBOC library screens are limited to ex vivo applications (e.g., on-cell selection), and therefore do not ensure tissue selectivity in vivo and fail to fully capture the mechanical processes that govern biodistribution. While these strategies have allowed progress in the discovery of CTPs, there is a critical need for a technique that can reliably generate D-CTPs in a form that functions efficiently within a living animal. We envisaged an in vivo selection of a D-peptide library as a feasible strategy to identify D-CTPs.

To discover an erythrocyte-binding D-CTP, we performed a selection on a randomized D-peptide library in live mice (Fig. 1a). A 10^6^-member 10-mer D-peptide library was synthesized with C-terminal amides to enable downstream spectral filtering from endogenous peptides, which bear C-terminal carboxylic acids. This accounts for a −0.98 Da mass difference on the library members, which can be reliably identified by mass spectrometry. Four mice were injected with amidated D-peptide library solution, after which they were euthanized. Blood and membrane-bound material was processed and prepared for mass spectrometry analysis (see Methods for detail). Nano-liquid chromatography-tandem mass spectrometry (nLC-MS/MS) analysis identified 128 putative blood-binding amidated 10-mer D-CTPs. A positional frequency analysis indicated a preference among these hits for a first-position D-phenylalanine and second-position D-tryptophan (Fig. 1b). The strongest amino acid preference was for a fourth position D-proline, which occurred in over half of library sequences identified from our randomized peptide library. Several sequences with varying resemblance to the consensus motif were individually synthesized as dye-labeled peptides. Reasoning that because of their abundance erythrocytes were the likely binding partner in a blood-based affinity selection, we used flow cytometry to determine the apparent binding constant of the putative binders to mouse erythrocytes (Fig. 1c and Supplementary Fig. 1). We observed that the highest affinity binders generally resembled the consensus motif. In particular, peptides with the consensus fourth position D-proline had measurable affinity for erythrocytes (though lyqpawfafk bound only weakly), and those without D-Pro4 did not. Of note, the highest affinity peptide measured, referred to hereafter as DQLR, also contained the consensus first-position D-phenylalanine and second-position D-tryptophan.

**Fig. 1.**
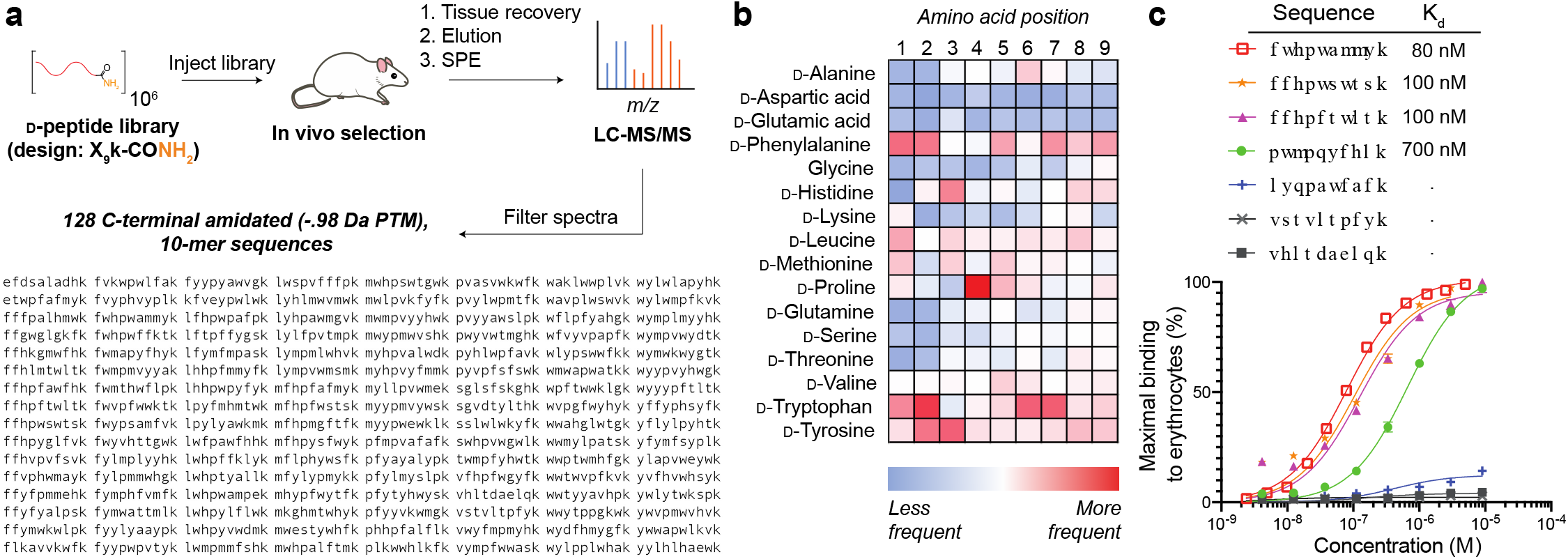
In vivo selection of a D-peptide library enables de novo discovery of nanomolar binders to murine erythrocytes. **a**, Description of our in vivo selection strategy to discover tissue-targeting peptides from a randomized 10^6^-member D-peptide library, and library design-matching peptide sequences identified from selection in four mice (10-mer sequences with C-terminal amidation). **b**, Positional frequency analysis of identified sequences indicates a preference for first-position d-phenylalanine, second position d-tryptophan, and fourth position d-proline. **c**, AlexaFluor488-labeled selection-derived peptides bind mouse erythrocytes with varying affinity, as determined by flow cytometry (n = 3; data presented as mean ± s.e.m.).

We established the sequence specificity of the DQLR-erythrocyte binding interaction. We compared the binding affinity of a DQLR-GFP conjugate to a DQLR-AlexaFluor488 conjugate, and found that both constructs retained their affinity for erythrocytes (Fig. 2a). DQLR-AlexaFluor488 bound to erythrocytes with an 80 nM dissociation constant, while DQLR-GFP had a 370 nM dissociation constant. We also examined the binding of a scrambled sequence of the DQLR peptide and the L-enantiomer of DQLR and found that neither had affinity for erythrocytes. Taken together, these findings suggest that the association of DQLR to erythrocytes is driven by the amino acid sequence and not by the reporter molecules used for detection. Moreover, the sequence- and stereo-specificity of DQLR binding likely involves molecular recognition, and not solely non-specific interactions (i.e., “greasiness”).

**Fig. 2.**
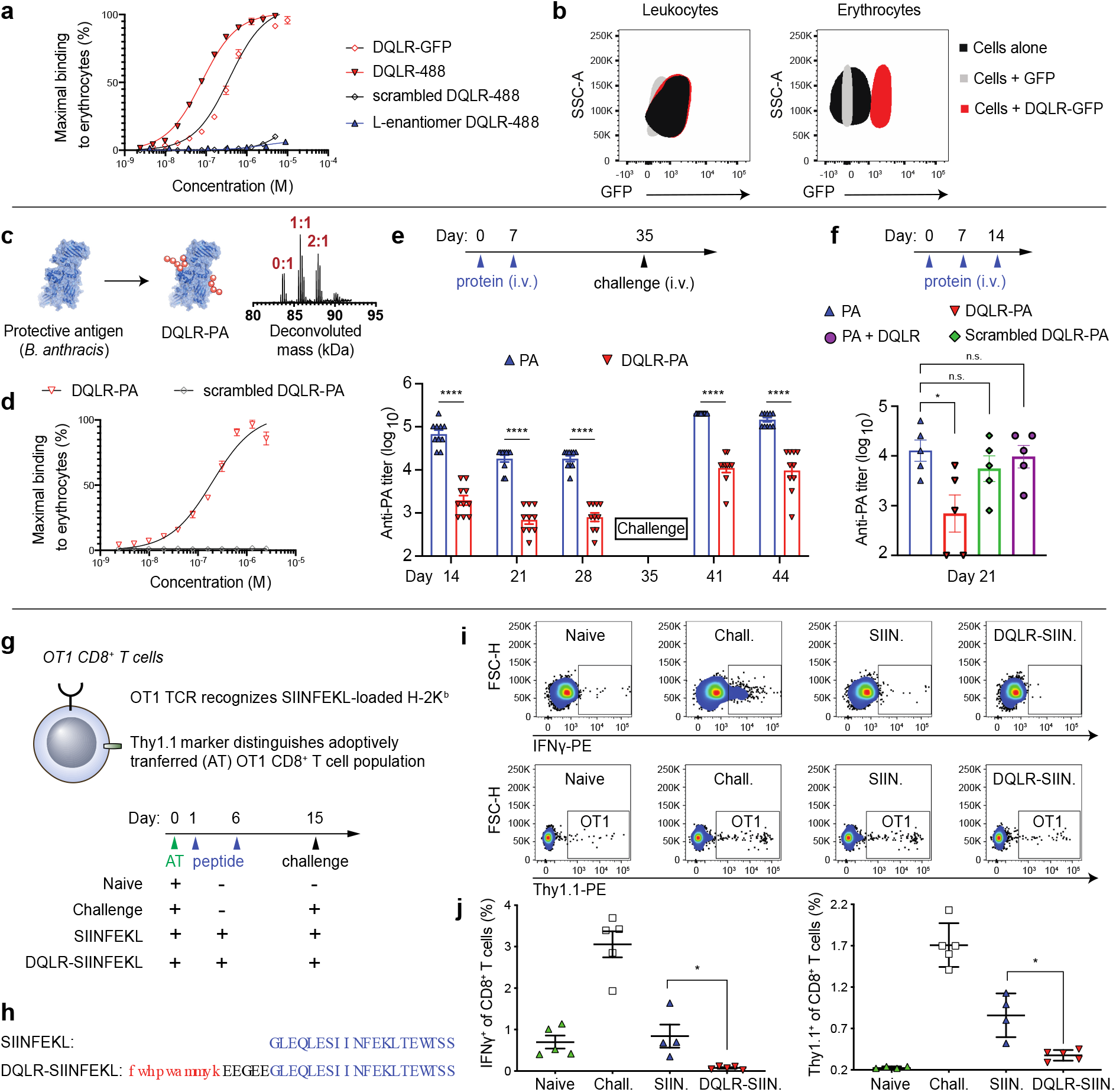
DQLR binds murine erythrocytes and its conjugates with peptide or protein antigens promote antigenspecific anti-inflammatory responses. **a**, AlexaFluor488-labeled DQLR and GFP-labeled DQLR bind mouse erythrocytes, whereas scrambled DQLR and L-enantiomer of DQLR do not, as determined by flow cytometry (n = 3; data presented as mean ± s.e.m.). **b**, GFP-labeled DQLR binds specifically to erythrocytes and not to splenic leukocytes, as determined by flow cytometry. **c**, Diagram and LC-MS deconvolution of DQLR-labeled mPA (K563C, N682A, D683A; DQLR-PA), with peptide: protein ratios in red. **d**, DQLR-PA binds to mouse erythrocytes, whereas scrambled DQLR-PA does not, as determined by flow cytometry (n = 3; data presented as mean ± s.e.m.). **e**, BALB/c mice have a decreased anti-PA antibody responses as a result of injection of DQLR-PA as compared to PA, even following a PA challenge to both groups (n = 10; data presented as mean ± s.e.m.). **f**, This decreased response is not observed when PA and DQLR are administered separately, or when scrambled DQLR-PA is administered (n = 5; data presented as mean ± s.e.m.). **g**, Diagram and injection scheme to determine SIINFEKL-specific antigen response. **h**, Sequences of SIINFEKL peptides injected. **i and j**, Mice administered DQLR-SIINFEKL have decreased pro-inflammatory T cell responses to SIINFEKL as determined by ex vivo SIINFEKL stimulation of peripheral blood-derived IFNγ^+^ CD8^+^ T cells, and measurement of draining inguinal lymph node-derived SIINFEKL-H-2K^b^-reactive OT1 CD8^+^ T cells (n = 4 for SIINFEKL group, n = 5 for other groups; data presented as mean ± s.e.m.).

We investigated the selectivity of the DQLR-erythrocyte binding interaction. We compared the binding of GFP and DQLR-GFP to mouse erythrocytes and leukocytes (Fig. 2b). To splenic leukocytes, GFP and DQLR-GFP displayed comparable binding at 1 μM concentration, slightly above the fluorescence of leukocytes alone. In contrast, DQLR-GFP bound erythrocytes above background levels of autofluorescence, whereas GFP did not. Taken together, this line of evidence suggests that DQLR binds erythrocytes selectively, and does not contribute to the weak association of GFP to leukocytes.

We determined the ability of DQLR to direct a full-length protein to erythrocytes and mediate an antigen-specific anti-inflammatory response. Protective antigen (PA) derived from *Bacillus anthracis* is a component of anthrax toxin, and is an immunogenic protein of therapeutic interest^13–16^. We prepared wild-type PA and a mutant form of PA (mPA, K563C, N682A, D683A) that does not bind to its receptors and bears a single solvent-exposed cysteine residue. mPA was labeled with maleimide-biotin, then non-specifically labeled with a heterobifunctional linker with NHS-ester and maleimide functionalities, and finally labeled this construct with DQLR peptides containing terminal cysteine residues (DQLR-SH), yielding biotinylated DQLR-PA with a peptide-protein ratio ranging from 0:1 to 2:1 as determined by LC-MS (Fig. 2c; median of 1:1). We confirmed by flow cytometry (using streptavidin-phycoerythrin) that DQLR-PA bound to erythrocytes, whereas scrambled DQLR-PA did not (Fig. 2d). Mice were given either PA or DQLR-PA and then challenged with PA (Fig. 2e). Anti-PA antibody titers determined before and after the challenge showed a durable, significant decrease in the anti-PA antibody response in the DQLR-PA group. Administration of scrambled DQLR-PA or a mixture of DQLR and PA did not significantly decrease the anti-PA response as compared to a animals that received only PA, indicating that covalent linkage to DQLR is required (Fig. 2f). This outcome suggests that physical association of antigens with erythrocytes has a tolerogenic effect, achievable by conjugating a protein antigen to the erythrocyte-binding D-peptide DQLR.

The erythrocyte-targeting D-peptide DQLR can also promote an anti-inflammatory response in the context of a peptide antigen. We used the peptide SIINFEKL, derived from full-length ovalbumin (OVA) protein (Fig. 2g). Mice received OT1 Thy1.1^+^ CD8^+^ T cells, derived from TCR transgenic mice specific for SIINFEKL-loaded H-2K^b^ complexes. Mice were then given either DQLR-SIINFEKL peptide or SIINFEKL peptide alone (Fig. 2h). Mice were later challenged via intradermal injection of full-length OVA and LPS. Mice that received OT1 Thy1.1^+^ CD8^+^ T cells as well as the OVA challenge, but were pretreated with PBS instead of the various peptides, served as controls. On day 19, peripheral blood leukocytes were stimulated ex vivo with SIINFEKL peptide, and the percentage of intracellular interferon-gamma positive (IFNγ^+^) CD8^+^ T cells was determined by flow cytometry. On day 21, draining inguinal lymph nodes were harvested and the percentage of Thy1.1^+^ CD8^+^ T cells was determined. Compared to mice that received SIINFEKL peptide alone, mice that received DQLR-SIINFEKL had significantly fewer IFNγ^+^ CD8^+^ T cells and SIINFEKL-H-2K^b^ −reactive OT1 CD8^+^ T cells (Fig. 2i, 2j, Supplementary Fig. 2, and Supplementary Fig. 3). Taken together, these results suggest that the DQLR-SIINFEKL peptide drove deletion of the OT1 T cell population and promoted an antiinflammatory response. Of note, under these experimental conditions, a conjugate of SIINFEKL to a previously reported erythrocyte-binding L-peptide (ERY1-SIINFEKL) did not significantly alter the level of IFNγ^+^ CD8^+^ T cells or OT1 CD8^+^ T cells, as compared to SIINFEKL alone (Supplementary Fig. 4). These outcomes support the utility of our in vivo selection approach toward de novo discovery of erythrocyte-selective D-peptides.

The inherent susceptibility to proteolysis of tissue-targeting L-peptides in vivo hampers their discovery and application. Our proof-of-concept study of a D-peptide library selected in live mice provides a novel D-peptide ligand that binds to mouse erythrocytes. When administered as an antigen-CTP conjugate, these molecules have anti-inflammatory effects. This in vivo selection technique marries the benefits of customizable synthetic peptide libraries to a high fidelity ‘in solution’ selection^17,18^. We envision that this platform can be used to discover synthetic CTPs that recognize targets beyond erythrocytes, including specific cells, tissues, and organs of therapeutic interest, including tumors. We also believe this method could be used to investigate structure-function relationships between the properties of synthetic peptides (e.g., stereochemistry, non-canonical functional groups, synthetic peptide structures and supramolecular configurations) and the complex biological and physical features in animal models. Moreover, DQLR-antigen conjugates merit further investigation as a therapeutic route for inflammatory and autoimmune disorders driven by key antigens and autoantigens.

## Methods

### Peptide synthesis and purification

Automated fast-flow synthesis of peptide-^α^carboxamides, as previously described^19^, was carried out using H-Rink amide-ChemMatrix resin (0.45 mmol/g, 0.1 mmol scale) at 90 °C. For manual, batch amino acid coupling and modification, fritted syringes (Torviq) containing peptide resin were washed with DMF. Fmoc-protected amino acids (5 eq) in HATU solution (0.38M, 4.5 eq) were activated with DIEA (15 eq) and added to the resin bed. Resin was incubated at r.t. for 20 min, then washed three times with DMF. Fmoc protecting group was removed using 20% piperidine in DMF. SIINFEKL peptide used in OT1 experiment and for ex vivo stimulation was purchased from Genscript.

Split-and-pool synthesis of the peptide library was carried out on 130 μm TentaGel resin (0.26 mmol/g). Splits were performed in 18 plastic fritted syringes on a manifold. Couplings were carried out using solutions of Fmoc-protected amino acids, HATU (0.38M in DMF; 0.9 eq relative to amino acid), and DIEA (1.1 eq for histidine; 3 eq for all other amino acids). Resin was incubated at r.t. for 20 min, then washed three times with DMF. Fmoc protecting group was removed using 20% piperidine in DMF. Resin was washed again with DMF.

Peptide cleavage from solid support and global deprotection were carried by adding a solution of 94% (v/v) trifluoroacetic acid (TFA), 2.5% (v/v) ethanedithiol, 2.5% (v/v) water, and 1.0% (v/v) triisopropylsilane to resin, and incubating at r.t. for 2 h. Peptide was precipitated and triturated three times using diethyl ether. Crude peptide was dissolved in 50/50 water/acetonitrile (0.1% TFA) and lyophilized.

Peptides were purified by mass-directed semi-preparative reversed phase HPLC (Agilent). Solvent A was water with 0.1% TFA and solvent B was acetonitrile with 0.1% TFA. Peptides were generally purified with a 0.5% B/min gradient on a C3 SB Zorbax column (9.4 x 250 mm, 5 μm). Fractions were analyzed by LC-MS and the purest fractions were pooled.

### In vivo selection

Library stock solution (7.4 mM) in PBS was filtered using a 0.2 μm SPIN-X filter (Corning) and pH was adjusted to 7. Library solution was administered by tail-vein to C57Bl/6J mice (Taconic). Mice were female and 27 weeks old. The exact volume administered was 170-200 μL. After 10 minutes, mice were euthanized by asphyxiation. Subsequently, a cardiac puncture was performed to retrieve as much blood as possible, and blood was collected into labeled EDTA-coated collection vials. The exact quantity of blood collected was 400-600 μL. Vials were stored on ice for roughly 30 minutes before tissue processing. For processing, blood samples (from in vivo selection) were transferred to a 1.6 mL microcentrifuge tubes. Blood was washed once with PBS, then centrifuged at 500 g for 3 min. Blood was then washed with RBC Lysis Buffer (Stem Cell Technologies), then centrifuged at 16 000 g for 10 min. This was repeated twice. Cell pellet was resuspended in 50/50 water/acetonitrile (0.1% TFA), incubated at r.t. for 10 min, then centrifuged at 16 000 g for 10 min. The supernatant was transferred to a new microcentrifuge tube and lyophilized overnight. Lyophilized material was resuspended in 95/5 water/acetonitrile (0.1% TFA) containing 6 M guanidine-hydrochloride. Suspension was centrifuged at 16 000 g for 10 min, and the supernatant was transferred to a new microcentrifuge tube. This was repeated until the pellet formed by centrifugation was no longer visible (~10 rounds of centrifugation). The final supernatant was purified by solid-phase extraction using a 100 μL bed C18 tip, according to the manufacturer’s protocol (Pierce). The eluate was lyophilized in a 200 μL strip tube overnight. Lyophilized material was resuspended in water (0.1% TFA) and centrifuged at 16 000 g for 5 minutes. The supernatant was analyzed by nLC-MS/MS.

### Preparation of proteins and protein conjugates

GFP was purified from *E. coli. E. coli* expressing G3-GFP were grown at 37 °C in LB broth to an OD_600 nm_ of 0.6, then grown overnight at 30 °C. Bacteria were centrifuged at 8 000 g at °C and stored at −80 °C overnight. The following day, pellets were thawed on ice and homogenized by magnetic stirring and using a sonication probe (Branson). Lysate was centrifuged at 8 000 g at 4 °C, and the supernatant was purified using an AKTA Pure FPLC equipped with a HisTrap FF crude 5 mL column (GE Healthcare). Solvent A was 20 mM tris pH 7.5. Solvent B was 20 mM tris, 300 mM imidazole pH 7.5. The gradient used was 0-50% B over 20 column volumes. Fraction purity was assessed by SDS-PAGE. The purest fractions were collected, filtered using a 0.2 μm filter, snap frozen in liquid nitrogen, and stored at −80 °C.

PA and mPA were prepared from *E. coli. E. coli* pellets containing periplasm-localized wild-type PA and mPA (K563C, N682A, D683A) were obtained from a protein expression facility and purified using an osmotic shock method. A pellet was thawed in sucrose buffer [20% sucrose (w/v), 20 mM tris, 1 mM EDTA, pH 8] and homogenized by magnetic stirring at °C. The cell homogenate was centrifuged at 8 000 g for 20 min at 4 °C. The supernatant was discarded, and the pellet was resuspended in 5 mM magnesium sulfate buffer and homogenized. The homogenate was stirred for 30 min at 4 °C. The homogenate was centrifuged as before and the supernatant was purified on an AKTA Pure FPLC equipped with a HiTrap Q 5 mL anion exchange column (GE Healthcare). Solvent A was 20 mM tris pH 8.5. Solvent B was 20 mM tris, 1 M sodium chloride pH 8.5. The gradient used was 0-25% B over 11 column volumes. Fraction purity was assessed by SDS-PAGE. The purest fractions were collected. These were further purified by size exclusion chromatography on a HiLoad 26/600 Superdex 200 pg column, using an isocratic 1.5 column volume elution employing 20 mM tris, 150 mM sodium chloride pH 7.5 buffer. The purest fractions, as determined by SDS-PAGE, were filtered using a 0.2 μm filter, snap frozen in liquid nitrogen, and stored at −80 °C.

DQLR-PA and scrambled DQLR-PA were prepared using non-specific labeling chemistry. mPA (K563C, N682A, D683A) was modified with 10 equivalents of biotin-maleimide (Sigma Aldrich) at r.t. for 30 min. Biotinylated mPA was then modified with 10 equivalents of sulfosuccinimidyl 4-(N-maleimidomethyl)cyclohexane-1-carboxylate (sulfo-SMCC; Thermo Fisher Scientific) at r.t. for 1 h. Finally, SMCC-modified biotinylated mPA was modified with 10 equivalents of DQLR-SH or scrambled DQLR-SH at r.t. for 1 h. After each reaction, protein was buffer exchanged three times into new buffer to remove unreacted small molecules and peptides. The final protein-peptide conjugates were filtered using 0.2 μm filters, snap frozen in liquid nitrogen, and stored at −80 °C.

DQLR-GFP was prepared by sortase ligation. DQLR-LPXTG (1 mM) was enzymatically ligated to G3-GFP (100 μM) using pentamutant sortase^20^ (5 μM) in sortase buffer (10 mM calcium chloride, 20 mM tris, 150 mM sodium chloride pH 7.5) at r.t. for 1 h. The reaction was then purified on an AKTA Pure FPLC equipped with a HiLoad 26/600 Superdex 200 pg column, using an isocratic 1.5 column volume elution employing 20 mM tris, 150 mM sodium chloride pH 7.5 buffer. The purest fractions, as determined by SDS-PAGE, were filtered using a 0.2 μm filter, snap frozen in liquid nitrogen, and stored at −80 °C.

### Mass spectrometry

LC-MS analysis was performed on a 6520 Electrospray Ionization Quadrupole Time-of-Flight (ESI-QTOF) LC-MS (Agilent) equipped with an Agilent C3 Zorbax column (300SB C3, 2.1 x 150 mm, 5 μm). Solvent A was water with 0.1% formic acid (FA) and solvent B was acetonitrile with 0.1% FA. The method used was 1% B for 2 min, 1-61% B over 9 min, 61-99% B over 1 min, and 3 min post-time at 1% B. Data was processed using Agilent MassHunter.

nLC-MS/MS was performed on an EASY-nLC 1200 (Thermo Fisher Scientific) nano-liquid chromatography handling system equipped with a PepMap RSLC C18 column (15 cm x 50 μm, 2 μm; Thermo Fisher Scientific) and a C18 nanoViper Trap Column (20 mm x 75 μm, 3 μm, 100 Å; Thermo Fisher Scientific), connected to an Orbitrap Fusion Lumos Tribrid Mass Spectrometer (Thermo Fisher Scientific). Solvent A was water (0.1% FA) and solvent B was 80% acetonitrile, 20% water (0.1% FA). The nano-LC gradient was 1% solvent B in solvent A ramping linearly to 41% B in A over 120 min (40 °C, flow rate of 300 nL/min). Positive ion spray voltage was 2200 V. Orbitrap was used for primary MS, with the following parameters: resolution = 120,000; quadrupole isolation; scan range = 200-1400 m/z; RF lens = 30%; AGC target = 1 x 10^6^; maximum injection time = 100 ms; 1 microscan. Acquisition of MS/MS spectra was done was performed in the Orbitrap (resolution = 30,000; quadrupole isolation; isolation window = 1.3 m/z; AGC target = 2 x 10^4^; maximum injection time = 100 ms; 1 microscan) in a data-dependent manner: a precursor was excluded for 30 s if it was detected greater than three times within 30 s (mass tolerance: 10.00 ppm); peptides were selected by monoisotopic precursor selection; intensity threshold was 5 x 10^4^; the 2–10 charge states were selected; the precursor selection range was 200-1400 m/z. The top 15 most intense precursors matching these criteria were subjected to subsequent fragmentation. Collision-induced dissociation (CID), higher-energy collisional dissociation (HCD), and electron-transfer/higher-energy collisional dissociation (EThcD) were used for MS/MS acquisition. Precursors with charge states greater than 2 were subjected to all fragmentation modes; precursors with charge states of 2 were subjected to CID and HCD only. A collision energy of 30% was used for CID. A collision energy of 25% was used for HCD. A supplemental activation collision energy of 25% was used for EThcD.

### Flow cytometry

Cells were analyzed on a FACS Canto II flow cytometer or FACS Fortessa flow cytometer (BD Biosciences). Flow data was processed in FlowJo (Tree Star). Cells (excluding erythrocytes) were stained with a fixable live/dead aqua zombie (Biolegend) for 15 minutes. The surface receptors were stained for 30 min on ice. For intracellular staining experiments, cells were fixed and permeabilized at this point using BD Cytofix/Cytoperm, then stained again. The following staining reagents were used: anti-CD45.2-BUV395 (BD Biosciences), anti-Thy1.1-PE (OT1 marker; Biolegend), anti-IFNγ-AF488 (Biolegend), anti-CD8-APC (Biolegend) streptavidin-PE (Biolegend), GFP, DQLR-GFP, biotinylated DQLR-PA, and biotinylated scrambled DQLR-PA. At least 1 000 000 events were collected for each experimental replicate, except for binding experiments in which at least 1 000 gated events were collected.

### Full-length protein immunogenicity

For the DQLR-PA versus PA experiment, ten BALB/c mice per group (female, 6-8 weeks old; Taconic) were anesthetized by isoflurane and administered 0.5 nmol of 0.2 μm-filtered protein in sterile PBS by retro-orbital injection, on “protein” days. All mice were challenged with 0.1 nmol wild-type PA on the “challenge” day. On bleed days, mice were bled via retro-orbital route (roughly 50 μL collected per bleed). Serum was collected and stored at −80 °C until ELISA was performed.

For the DQLR-PA versus PA versus DQLR + PA versus scrambled DQLR-PA experiment, five BALB/c mice per group (female, 8-10 weeks old; Taconic) were anesthetized by isoflurane and administered 0.5 nmol of 0.2 μm filtered protein in sterile PBS by tail-vein injection on “protein” days. On bleed days, mice were bled via submandibular routes (roughly 50 μL collected per bleed). Serum was collected and stored at −80 °C until ELISA was performed.

Anti-protein antibody titer was determined by ELISA. Medisorp 96-well plates (Thermo Fisher Scientific) were coated overnight at 4 °C with 10 μg/mL protective antigen in PBS. Plates were washed with PBST (PBS with 0.1% tween-20) and blocked with 5% FBS in PBS for 2 h at r.t. Subsequently, plates were washed with PBST and incubated for 2 h at r.t. with serum dilutions including a control well which was not antigen-coated. Plates were then washed and incubated with anti-mouse-horseradish peroxidase (1:5000 dilution in PBS; Thermo Fisher Scientific) and visualized with 1-Step Ultra-TMB (Thermo Fisher Scientific). The relative antibody titer was calculated as the highest dilution with greater than twice the OD_450 nm_ of the control well.

### Peptide immunogenicity

Peptides were tested for their ability to deplete CD8^+^ T cells toward SIINFEKL and a pro-inflammatory T cell responses using previously reported methods^2^. Briefly, 300,000 OT1 CD8^+^ T cells were administered intravenously (i.v.) via tail-vein injection into 6-8 week old C57Bl6/J mice (n = 4 or 5 per group as shown; Taconic). 60 μg of peptide was then injected i.v. on days 1 and 6 retro-orbitally. On day 15, mice were challenged with an intradermal injection of OVA and LPS, as previously described. 19 days post adoptive transfer, OT1 CD8^+^ T cells from the peripheral blood were quantified, as well as their activation by SIINFEKL by flow cytometric analysis. On day 21, the inguinal lymph nodes were harvested and OT1 CD8^+^ T cells were quantified by flow cytometry.

### Statistical analysis

Statistical analysis and graphing was performed using Prism (GraphPad). The listed replicates for each experiment indicates the number of distinct samples measured for a given assay. Significance for PA versus DQLR-PA was determined using a one-way ANOVA with a Sidak’s multiple comparisons test (\*\*\*\**P* < 0.0001). Significance for PA versus DQLR-PA versus DQLR + PA versus scrambled DQLR-PA was determined using a one-way ANOVA with a Dunnett’s multiple comparisons test (\**P* < 0.05). Significance for the OT1 experiments was determined using Mann-Whitney tests (\**P* < 0.05).

## Ethical compliance

All experiments described were performed in compliance with the Massachusetts Institute of Technology Committee on Animal Care (Institutional Animal Care and Use Committee).

## Data availability

All data needed to support the conclusions in the paper are present in the paper and supplemental information. Additional data is available upon request.

## Acknowledgements

Financial support for this work was provided by the MIT/NIGMS Biotechnology Training Program (T32 GM008334) to A.R.L. and A.J.Q. The authors would like to thank the MIT Preclinical Modeling, Imaging, and Testing core, and in particular Aurora Connor for her insight and contributions; the MIT Division of Comparative Medicine and in particular Paul Chamberlain and Elizabeth Horrigan for their assistance with studies and training; the NERCE facility (grant U54 AI057159) and Erica Gardner for protective antigen expression; Arisa Shimada for preparation of the peptide library reagent; Nina Hartrampf for assistance with automated peptide synthesis; and Nicholas Truex, Chi Zhang, Ronald T. Raines, and Alex K. Shalek, for support and helpful discussions.

## Author Contributions

A.R.L., G.Z., C.B., A.J.Q., R.J.C., H.P., and B.L.P. designed research; A.R.L., G.Z., C.B., A.J.Q., N.P., C.K.S., and D.G. performed experiments, with support from H.P., D.J.I., and B.L.P.; A.R.L., G.Z., C.B., A.J.Q., N.P., C.K.S., A.L., R.J.C., H.P., and B.L.P. analyzed data; A.R.L., C.B., and N.P. generated figures; and A.R.L. wrote the paper with input from all authors.

## Corresponding author

Correspondence to Bradley L. Pentelute.

## References

1. Pishesha, N. et al. Proc. Natl. Acad. Sci. 114, 3157–3162 (2017).

2. Kontos, S., Kourtis, I. C., Dane, K. Y. & Hubbell, J. A. Proc. Natl. Acad. Sci. 110, E60–E68 (2013).

3. Lorentz, K. M., Kontos, S., Diaceri, G., Henry, H. & Hubbell, J. A. Sci. Adv. 1, e1500112 (2015).

4. Serra, P. & Santamaria, P. Nat. Biotechnol. 37, 238–251 (2019).

5. Svensen, N., Walton, J. G. A. & Bradley, M. Trends Pharmacol. Sci. 33, 186–192 (2012).

6. Liu, R., Li, X., Xiao, W. & Lam, K. S. Adv. Drug Deliv. Rev. 110–111, 13–37 (2017).

7. Higgins, L. M. et al. Pharm. Res. 21, 695–705 (2004).

8. Ren, Y. et al. Mol. Pharm. 15, 592–601 (2018).

9. Ying, M. et al. ACS Appl. Mater. Interfaces 8, 13232–13241 (2016).

10. Wei, X. et al. Angew. Chemie Int. Ed. 54, 3023–3027 (2015).

11. Wei, X. et al. Mol. Pharm. 11, 3261–3268 (2014).

12. Yao, N. et al. J. Med. Chem. 52, 126–133 (2009).

13. Liu, S. et al. Proc. Natl. Acad. Sci. 113, E4079–E4087 (2016).

14. Loftis, A. R. et al. ChemBioChem n/a, (2020).

15. Gladstone, G. P. Br. J. Exp. Pathol. 27, 394–418 (1946).

16. Young, J. A. & Collier, R. J. Annu. Rev. Biochem. 76, 243–265 (2007).

17. Quartararo, A. J. et al. Nat. Commun. 11, 3183 (2020).

18. Pasqualini, R. & Ruoslahti, E. Nature 380, 364–366 (1996).

19. Hartrampf, N. et al. Science. 368, 980–987 (2020).

20. Chen, I., Dorr, B. M. & Liu, D. R. Proc. Natl. Acad. Sci. 108, 11399–11404 (2011)

