## Supplementary materials for "Erythrocyte-targeted immunomodulatory antigens enabled by in vivo selection of D-peptides"

**Affiliations:**

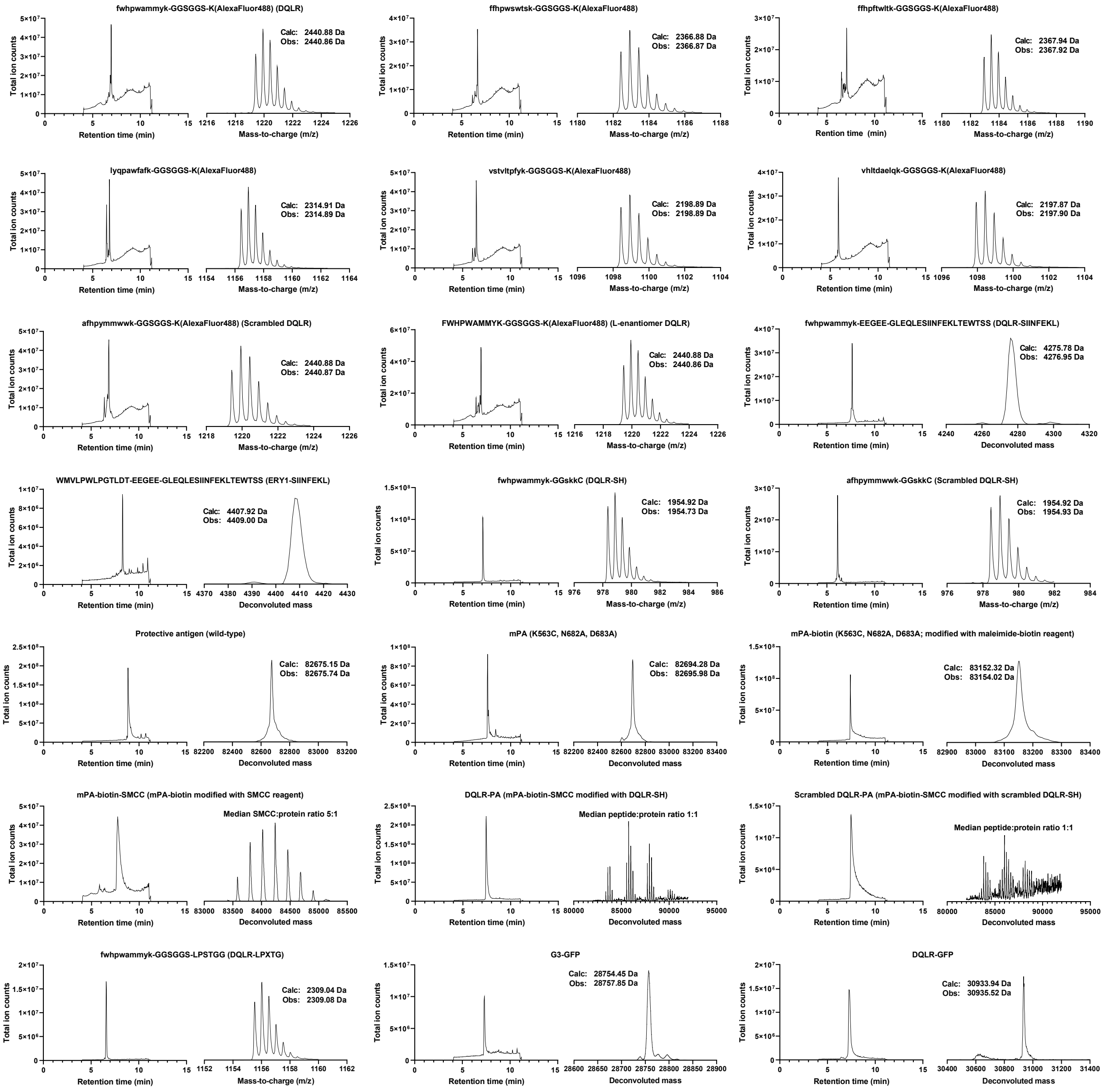


**Supplementary Figure 1. LC-MS analysis of peptide and protein constructs.** LC-MS analysis of peptide and protein constructs employed, including total ion chromatogram and either the extracted ion chromatogram or the mass deconvolution result used to calculate the observed mass.

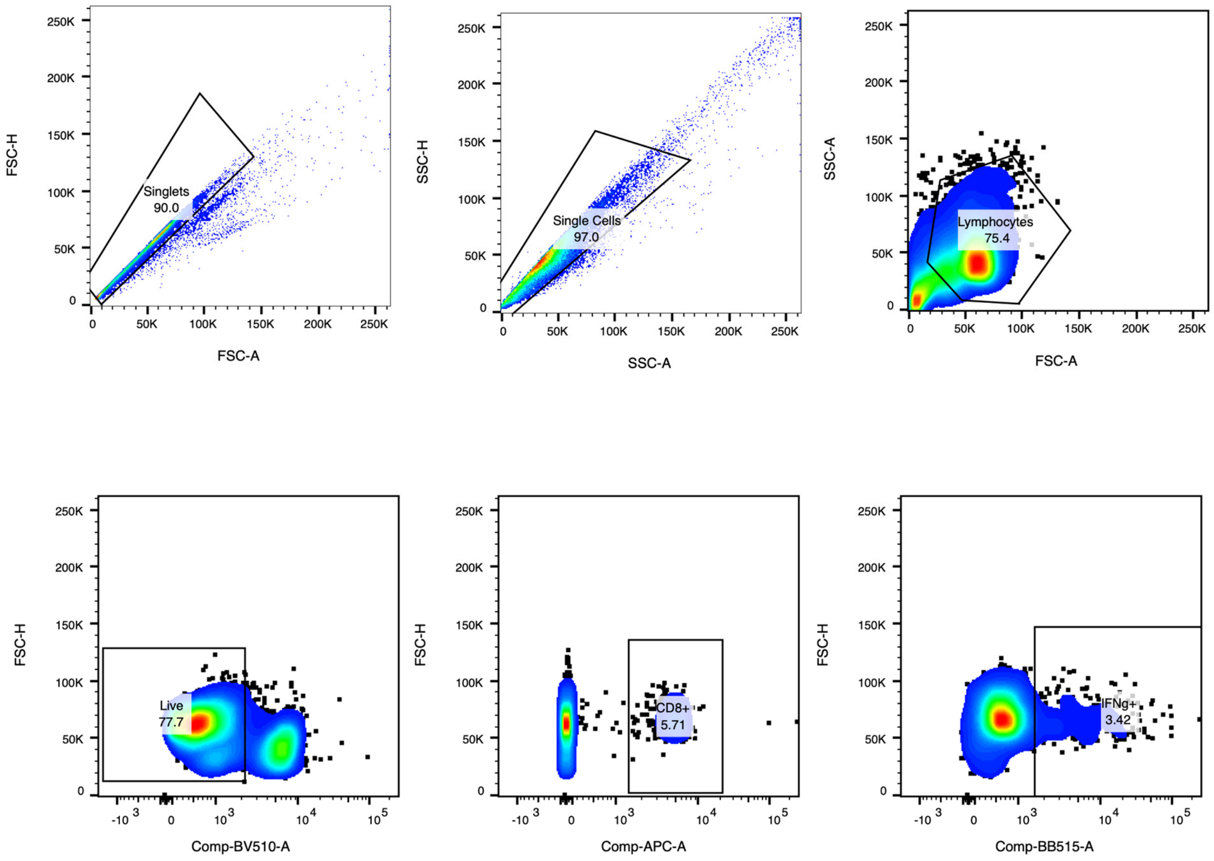


**Supplementary Figure 2. Gating strategy for IFNγ⁺ CD8⁺ T cells.** Flow cytometry gating strategy used to identify IFNγ⁺ CD8⁺ T cells following ex vivo stimulation with SIINFEKL peptide.

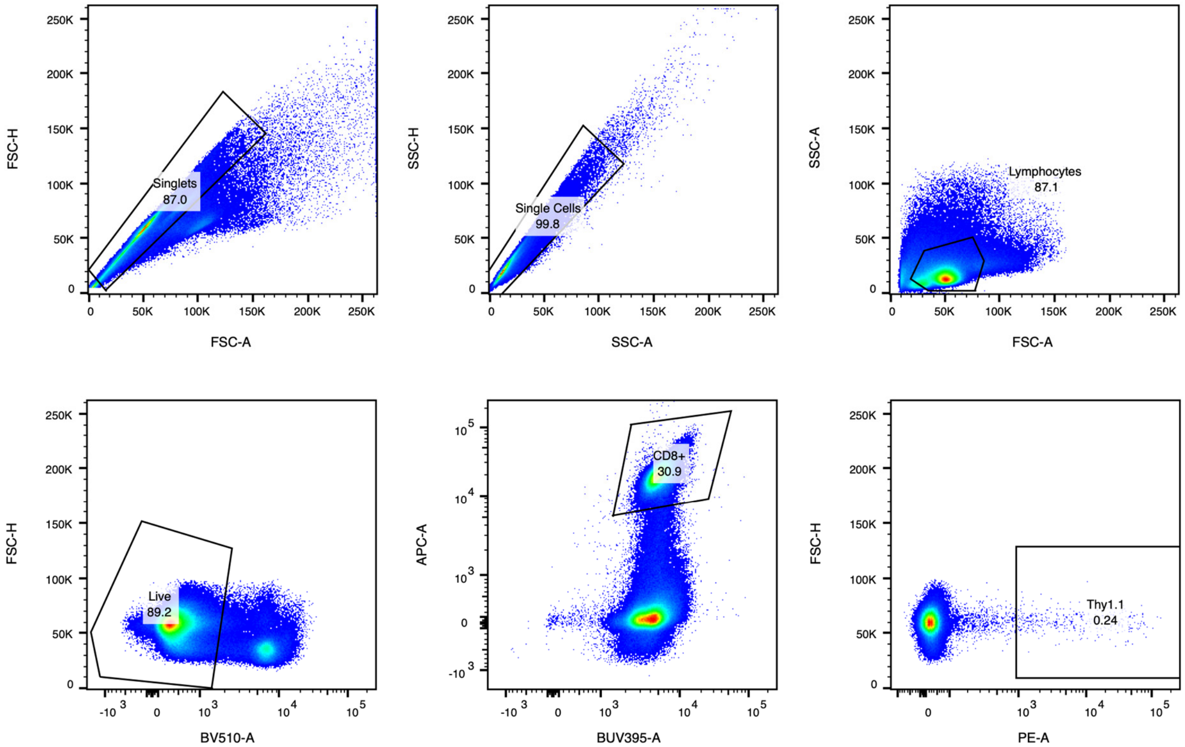


**Supplementary Figure 3. Gating strategy for Thy1.1⁺ CD8⁺ T cells.** Flow cytometry gating strategy used to identify circulating adoptively transferred OT1 CD8⁺ T cells.

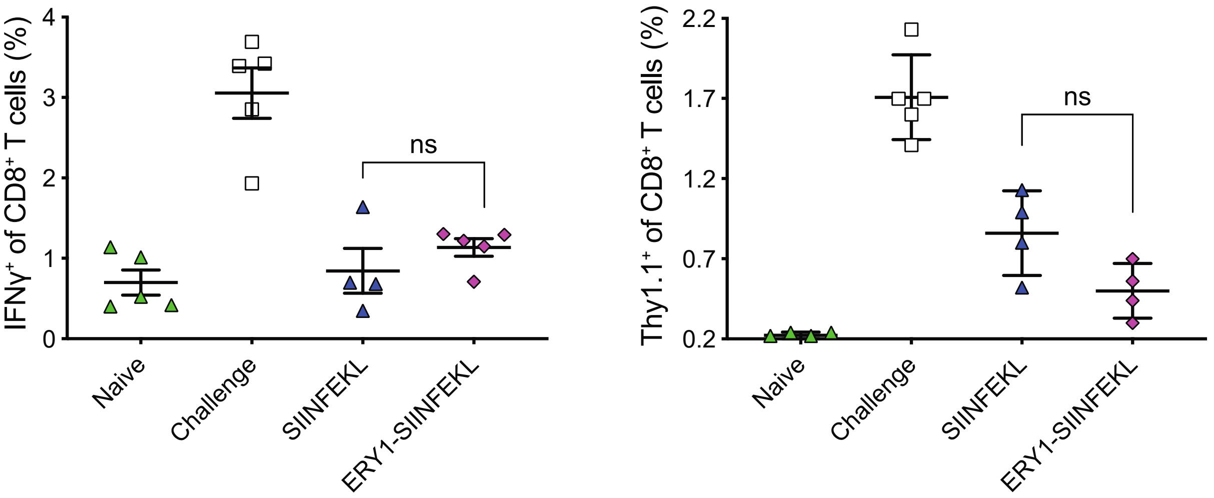


**Supplementary Figure 4. ERY1-peptide antigen conjugate does not significantly decrease the antigen-specific immune response to OVA.** Previously identified L-chirality erythrocyte-binding peptide ERY1 was prepared as a conjugate to SIINFEKL and administered to mice, which were adoptively transferred OT1 $CD8^+$ T cells and subsequently challenged with full-length OVA and LPS (as described for DQLR-SIINFEKL). The conjugate, ERY1-SIINFEKL, did not significantly decrease the inflammatory IFNγ response or the number of OT1 (i.e., $Thy1.1^+$) T cells, as compared to SIINFEKL alone (data presented as mean ± s.e.m.).
